# Small RNA Analysis of Virus-virus Interaction between Two Orthotospoviruses

**DOI:** 10.1101/2023.08.28.555202

**Authors:** Kaixi Zhao, Md Tariqul Islam, Nathan R. Johnson, Michael J. Axtell, Cristina Rosa

## Abstract

Mixed infections of plant viruses are commonly found in natural patho-systems and present a valuable opportunity to understand how multiple viruses can co-infect the same host. Tomato spotted wilt orthotospovirus (TSWV) and impatiens necrotic spot orthotospovirus (INSV) are present in the same geographic areas and are closely related. More mixed infections of TSWV and INSV have been reported in recent years, and the INSV host range has been reported to be increasing. In a previous study, we have isolated one strain of INSV and one of TSWV and showed that they have an antagonistic relationship in their vectors, but we were unable to determine, the underlying mechanisms governing their antagonism *in planta* and the contribution of the host to this. Here, we used small RNA sequencing to study TSWV-INSV antagonistic interaction and showed that INSV alters plant responses and the processing of TSWV.

## 1. Introduction

Viral synergy was first described in the 1950s with mixed infection of potato virus X (PVX), a potexvirus, and potato virus Y (PVY), a potyvirus. Simultaneous inoculation of those two viruses resulted in a much more severe disease than infections caused by either virus alone [1]. The beneficiary virus, which in this case was PVX, reached a higher titer compared to the titer seen in single infection, whereas PVY accumulation was not affected [2]. The mechanism behind this synergy was later demonstrated to be mediated by the potyvirus’ strong silencing suppressor protein P1/HC-Pro that enhanced the synergistic effect [3]. While the experiment performed in the above paper between PVX and PVY in *Nicotiana tabacum* showed 3-10 folds increase in PVX RNA replication intermediates [1,2], no change in viral titer or in host responses were observed in *N. benthamiana* plants during a synergistic interaction between the same two viruses [4,5]. This suggests that differences in host, virus strains, or experimental conditions could influence the outcome of virus interaction. Furthermore, how synergism works at the molecular level is still not clear [6]. A comparative transcriptional analysis revealed that during PVY and PVX synergism in *N. benthamiana* there is a difference in the expression level of oxidative stress related genes against α-dioxygenase-1 [7,8]. This is likely triggered by RNAi interference (RNAi) and is another indication that gene silencing could play a role in the outcome of co-infections.

Synergistic interaction happens also between tomato spotted wilt orthotospovirus (TSWV) and tomato chlorosis virus (ToCV, genus *Crinivirus*). Plant resistance against orthotospoviruses can be overcome by multiple infections, as when *Sw-5* resistant tomatoes are first infected with ToCV and then by a delayed inoculation (10 days later) with TSWV.

The authors speculated that ToCV could encode a repressor of the resistance response in *Sw-5* plants. Interestingly, simultaneous inoculation of a mixture of those two viruses results in no TSWV infection, as expected in orthotospovirus resistant plant genotypes [9]. Recent studies have shown that *Sw-5* resistance is based on a classical gene for gene interaction between the tomato coil-coil intracellular nucleotide binding leucine-rich repeat (CC-NLR) immune receptor *Sw-5b* and the orthotospovirus encoded movement protein NSm [10]. Still, the downstream pathway activated by this interaction that leads to hypersensitive response and programmed cell death, and thus to resistance, is unknown and could include a more generalized pathway such as RNAi.

Not all virus coinfections result in synergy, in fact antagonistic relationships have been described between unrelated potyviruses and potexviruses [11,12]; and between begomoviruses and tobamoviruses [13].

While, as indicated above, unrelated viruses with few exceptions interact with each other in a synergistic way, for related viruses, the interactions are mostly antagonistic. These antagonistic relationships, also called super-infection exclusion (SIE) [14], exclude a virus from tissues already infected by another virus and can be used as form of cross-protection in agriculture, where mild virus strains are used to ‘vaccinate’ plants against severe strains of the same virus [15](15). Super-infection exclusion has often been attributed to RNAi, but recent studies have suggested that some viruses can exclude highly similar or nearly identical viruses from the same cells by using a protein-based mechanism that acts on virus replication [16,17].

To complicate matters, many studies have now shown that the order of virus infection could change how viruses interact. Sequential inoculation of PVX and PVY yielded severe symptoms like the ones seen as consequence of their co-inoculation only when PVX was inoculated before PVY, while if PVY was inoculated before PVX symptoms were less severe and PVX titer was not increased as dramatically [1]. In a follow up study, Goodman and Ross (1974) found that the two viruses needed to be replicating in the same cells simultaneously to obtain the maximum benefit from their interaction [18]. In another example, Chávez-Calvillo and colleagues found antagonism when papaya ringspot virus (PRSV, genus *Potyvirus*) infection occurred after papaya mosaic virus (PapMV, genus *Potexvirus*) inoculation in papaya, and synergism when PRSV was inoculated 30 days before or together with PapMV [12]. Translation of PapMV RNA was not affected when the antagonistic interaction occurred, but the RNA accumulation of PRSV decreased and production of reactive oxygen species (ROS) and pathogenesis related protein PR-1increased by 2.2-folds during co-infection. These results suggest that PRSV infection could compromise the translation efficiency of non-PRSV mRNAs. Next generation sequencing (NGS) showed that the abundance of virus small interfering RNAs (vsiRNAs) was similar between synergistic and antagonistic interaction [12], suggesting that the RNAi pathway was not responsible for this difference.

TSWV and impatiens necrotic spot orthotospovirus (INSV) are orthotospoviruses that belong to the same phylogenetic ‘American clade’ [19,20]. Orthotospovirues share the same genome organization, with 3 genomic segments made of negative sense RNA encoding respectively for an RNA dependent RNA polymerase (on the large or L segment), a polyglycoprotein and a movement protein (on the medium or M segment), and a nucleocapsid protein and a silencing suppressor (on the small or S segment). Multiple protein coding regions, if they are on the same RNA segment, are separated by intergenic regions (IGRs) and all genomic segments start and end with untranslated regions (UTRs). The two TSWV and INSV isolates we used in this study [21,22] share less than 80% sequence similarity in the S region coding for the nucleocapsid protein, the sequence most used to define orthotospoviruses at the species level. Mixed infections of TSWV and INSV occur in agricultural crops and were first reported in tomato plants from Italy in 2000 [23]. In the US they were reported in tobacco in six counties in Georgia, Florida, South Carolina and Virginia in 2002 [24], where the incidence of co-infection in tobacco was up to 40% [25], and in North Carolina and Kentucky since 2003 [24]. Because INSV single infection in the study by Martinez-Ochoa was never detected in tobacco but only in the surrounding weeds, authors suggested that INSV could act as helper virus during mixed infection, indicating that TSWV and INSV have a synergistic and not antagonistic relationship during mixed infection [25]. On the contrary, in our studies that used two Pennsylvania isolates of the two viruses, INSV and TSWV showed an antagonistic relationship in their insect host, where INSV negatively impacted the ability of thrips to retain and transmit TSWV and suggesting that vectors could serve as bottleneck for the establishment and maintenance of TSWV and INSV mixed infection [26]. While our study did not find any significant difference between plant volatiles emitted during single and double infection, or between symptoms and infected plant appearance and size, thrips were able to distinguish between treatments, suggesting that plant respond differently when infected by one or two viruses [26]. Hypothetically, the result and establishment of a mixed infection could depend on the direct interaction of the two viruses (for instance, on direct competition for host resources), on plant responses to infection and on the propensity of the virus vectors two transmit one virus, the other of both simultaneously.

RNAi is the main mechanism used by plants to fight against viruses [15,27,28]. RNAi is triggered by the presence of (dsRNA) formed during viral replication or hairpin RNA produced by self-complementarity. These RNA forms elicit their processing by the plant Dicer-like (DCL) enzymes into small RNAs (sRNAs) of approximately 21–24 nt [29]. One of the two sRNAs strands is incorporated into the RNA-induced silencing complex (RISC) and used to guide Argonaute (AGO) to recognize, by sequence complementarity, the target RNA (in this case the virus RNA) and degrade it [30]. Secondary sRNAs are also produced in plants by the activity of a plant RNA-dependent RNA-polymerase and seem to be involved in magnification and spread of silencing [31]. On the other hand, viruses are known to produce proteins, called silencing suppressors, that modulate and down-regulate or interfere with different steps of the RNAi pathway [27] TSWV silencing suppressor NSs, for instance, has been shown to interfere with the binding of DCL to dsRNA and to siRNAs [32–34]. Next generation sequencing and analysis of sRNA profiles have been used to study plant virus interaction and as consequence to enhance our understanding of host responses towards virus infection [35–37]. Small RNA profiles of TSWV have been reported [38,39] but no INSV sRNA profile is available.

Based on the studies cited above, it is not clear if TSWV and INSV have a synergistic or antagonistic interaction, thus, we decided to delve more into this interaction in plants. Here we hypothesized that the plant host plays a role in TSWV and INSV co-infection outcomes and that a signature of this role would be retained in the sRNAs profile of co-infected plants. Determining if the host defense system (i.e., RNAi) responds similarly to TSWV and INSV in single or mixed infection could also provide novel orthotospovirus species or genus specific targets to use for disease management.

## 2. Materials and Methods

### 2.1. Viruses and their maintenance

*Nicotiana benthamiana* plants at 3-4 leaf stage and grown from seed were mechanically inoculated with TSWV isolate PA01[21] and/or INSV isolate UP01 [22], using virus-infected tissue from the originally infected hosts stored at -80 °C as source of inoculum. Inoculated plants were maintained in growth chamber at a temperature of 25 °C with a 16/8 h light/dark photoperiod and monitored for symptoms development.

### 2.2. Virus-virus interaction

Since at the time of this experiments no infectious clones of these two viruses were available, competition assays were performed using mechanical inoculation of infected plant extract. *N. benthamiana* plants were mechanically inoculated with 5 dilutions of a single virus inoculum and results were compared with the ones obtained by mechanically inoculating plants with the same 5 dilutions of inoculum spiked with an equal amount of the second virus. Inoculated plants were maintained in growth chamber at a temperature of 25 °C with a 16/8 h light/dark photoperiod and monitored for symptoms development.

### 2.3. Sequential inoculation for study of super infection exclusion

*N. benthamiana* plants were mechanically inoculated with INSV or TSWV infected inoculum on the left half of two leaves in each plant. Forty-eight hours after the first inoculation, the right half of the same two leaves was inoculated with equal amount (w/w) of the other virus. Systemic leaves were tested by ELISA (Agdia, Elkhart, IN, USA) following the manufacturer protocol for the presence of each virus 2 weeks after inoculation. The experiment was repeated twice.

### 2.4 Virus inoculations for sRNA sequencing

A batch of 10 *N. benthamiana* plants for each single infection and 20 plants for mixed-infection mechanically inoculated using the following scheme: Treatment 1: TSWV PA01 and healthy *N. benthamiana* (1:1, w/w); Treatment 2: INSV UP01 and healthy *N. benthamiana* (1:1, w/w); Treatment 3: mixture of TSWV PA01 and INSV UP01 (5:2, w/w) as the source of mixed-infection; and Treatment 4: mock-inoculated with healthy *N. benthamiana* leaves [40].

### 2.5. RNA extraction, qPCR and sRNA sequencing

Three systemically infected leaves from each plant were sampled and stored at −80 °C. Three leaf disks were sampled from those three systemically infected leaves (one leaf disk per leaf) for virus titer quantification by ELISA. Leaf tissue showing no statistically significant differences in virus titer at the intratreatment level were used for RNA extraction and library preparation. Three leaf disks (one leaf disk per leaf) of two plants in the same treatment group were pooled to generate one library for sRNA sequencing. Two libraries were performed for each treatment to generate biological replicates. Total RNA was extracted with the Quick-RNA miniprep kit (Zymoresearch, Irvine, CA, USA) following the manufacturer protocol. RNA was quantified by NanoDrop 2000 spectrophotometer (Thermo Scientific, Waltham, MA, USA).

Aliquots containing ∼500ng total RNA were sent to the Genomics Core Facility at the Pennsylvania State University for sequencing. RNA quantity was measured by Qubit (Thermo Fisher, Waltham, MA, USA) and quality was measured by Agilent Bioanalyzer (Santa Clara, CA, USA). Libraries were made using the TruSeq Small RNA Library Preparation Kit following the manufacture protocol (Illumina, San Diego, CA, USA) and then sequenced by Illumina NextSeq 550.

qPCRs on the RNA sent for sequencing were preformed using primers from [22] and iTaq Universal SYBR Green Supermix following the manufacture protocol (Bio-rad, Hercules, CA, USA).

### 2.6. Bioinformatic analysis

Small RNA reads were trimmed for adaptor sequences and mapped against the viral (NCBI accession numbers KT160280-KT160282 for TSWV and MH171172–MH171174 for INSV) [21,22] and <u>host</u> genomes (*N. benthamiana* genome version 0.5) [41] using Short-stack version 3.8.5 [42], allowing zero mismatches. Read size, distribution along the genomic segments and along single ORFs, polarity, 5′-nt enrichment and hotspot analyses were performed using <u>SAMtools</u> version 1.9 [43] and in-house Perl scripts, and imaged using Microsoft Excel® v. 10. Relative frequencies of 5’ terminal nucleotides and vsRNAs hotspot profiles were generated by MISIS 2.7 [44]. Differential expression of miRNA loci was analyzed using DESeq2 [45] with a log2 fold threshold of 1, and alpha of 0.1. Benja-mini-Hochberg procedure were used for multiple testing *P* values adjustment.

## 3. Results and Discussion

### 3.1. INSV show an antagonistic interaction against TSWV

TSWV and INSV were able to infect the same *N. benthamiana* plant when transmitted by mechanical inoculation. The number of plants infected in mixed infection differed from the number of plants infected by the single viruses, even when using the same inoculum and the same ratio of inoculum (Table 1), demonstrating that there is an interaction between the two viruses (Table 1). Disease incidence data showed 80-100% of the plants were infected by INSV in both single and mixed infection with various percent of inoculum. However, in case of TSWV, only 0-20% of the plants in mixed infection were found positive, whereas single infection of TSWV showed as high as 100% disease incidence with various percent of inoculum. This whole experiment has also been repeated with a more virulent TSWV isolate where higher virulence improved the chances for TSWV to infect plants in mixed infection (results not shown), however, INSV always demonstrated better fitness than TSWV.

**Table 1.** Virus-virus interaction experiment setup and percentage of infected plants from virus-virus interaction assays.

| Treatment 1a<br>(Control)% | INSV<br>Infected | Treatment 2a<br>(Mixed%) | INSV Infected | TSWV Infected | Mixed-infected |
| --- | --- | --- | --- | --- | --- |
| INSV(X) | 100% | INSV( X) + TSWV(X) | 100% | 40% | 40% |
| INSV(0.8X) | 100% | INSV(0.8X) + TSWV(X) | 100% | 20% | 20% |
| INSV(0.6X) | 80% | INSV(0.6X) + TSWV(X) | 100% | 20% | 20% |
| INSV(0.4X) | 100% | INSV(0.4X) + TSWV( X) | 80% | 60% | 40% |
| INSV(0.2X) | 60% | INSV(0.2X) + TSWV(X) | 0 | 60% | 0 |
| Treatment 1b<br>(Control)% | TSWV<br>Infected | Treatment 2b<br>(Mixed%) | TSWV Infected | INSV Infected | Mixed-infected |
| TSWV(X) | 100% | TSWV(X) + INSV(X) | 0 | 100% | 0 |
| TSWV(0.8X) | 100% | TSWV(0.8X) + INSV(X) | 20% | 100% | 20% |
| TSWV(0.6X) | 60% | TSWV(0.6X) + INSV(X) | 20% | 100% | 20% |
| TSWV(0.4X) | 20% | TSWV(0.4X) + INSV(X) | 20% | 100% | 20% |
| TSWV(0.2X) | 0 | TSWV(0.2X) + INSV(X) | 0 | 100% | 0 |

Nonetheless, INSV and TSWV had an antagonistic relationship, in fact the presence of the second virus decreased the titer of the first virus, in a dose dependent manner (Figure 1). INSV overall was shown to be a better competitor than TSWV, and the presence of TSWV in plants with mixed infection did not decrease INSV incidence but decreased its titer even in plants where TSWV was not detected by ELISA (Figure 1).

**Figure 1.**
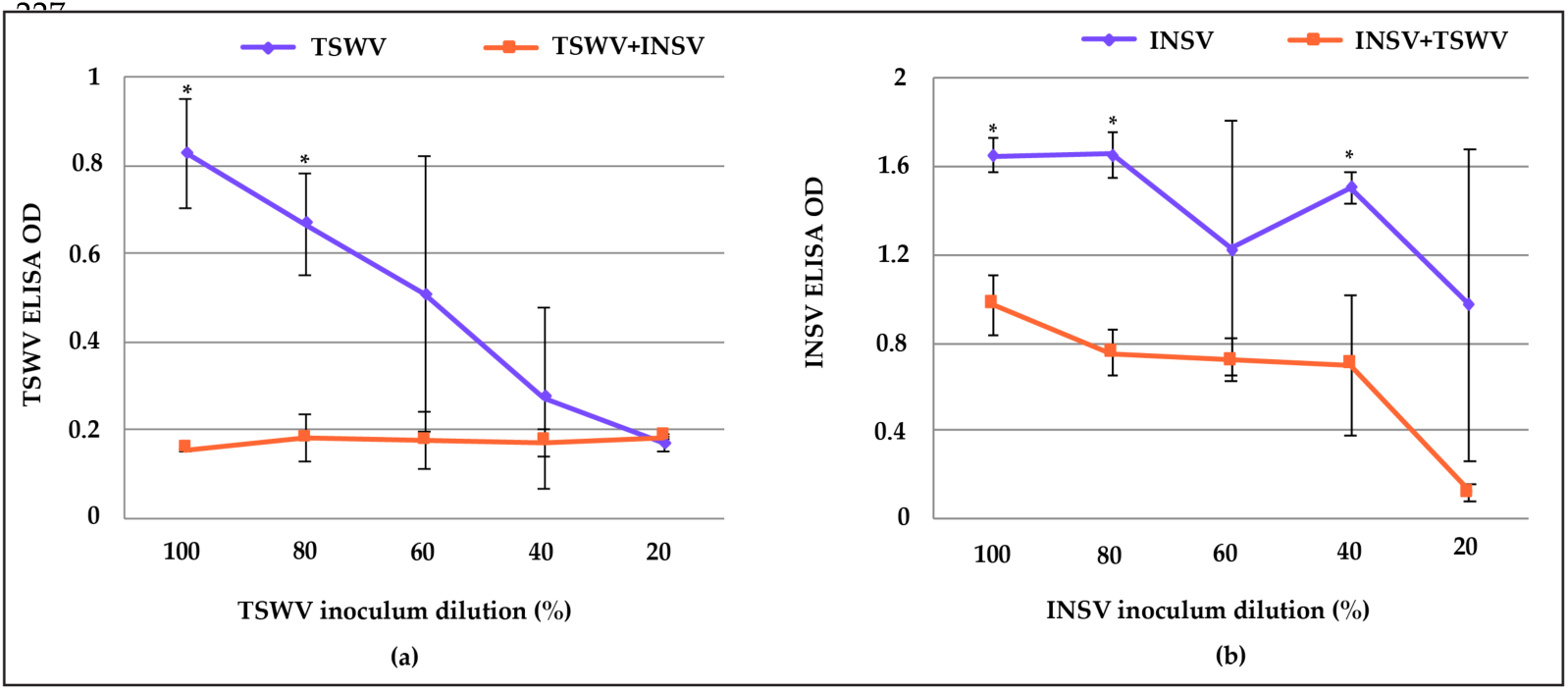
ELISA data showing antagonistic interaction of TSWV and INSV. X-axis indicates the percent of virus inoculum (TSWV or INSV infected leaf tissue in grinding buffer) and the remaining up to 100% was made by ground up healthy plant tissue. **(a)** TSWV titer was decreased by INSV in the mixed infection treatment (orange line) compared to single TSWV infection (violet line) and **(b)** INSV titer was decreased by the presence of TSWV in the inoculum (orange line), compared to single INSV infection (violet line). *N. benthamiana* plants were mechanically inoculated with 5 dilutions of a single virus and ELISA results were compared with the ones obtained by mechanically inoculating plants with the same 5 dilutions of inoculum spiked with an equal amount of the second virus. Error bar represents standard deviation, asterisks indicate difference from the single virus infection (P < 0.05) using two-sample t-test.n=5 for each treatment.

### 3.2. No cross protection between TSWV and INSV

When TSWV and INSV were inoculated sequentially, mixed infection could be detected in several plants. More specifically, when TSWV was inoculated before INSV, the overall mixed infection rate was 43.3% (n=30, 13 plants were infected with both, 12 plants were infected only with INSV, and 5 plants were infected only with TSWV). When INSV was inoculated first, the mixed infection rate was 29.4% (n=34, 10 plants were infected with both, 21 plants were infected with only INSV, and 3 plants were infected with only TSWV). Plants inoculated only with one virus were used as controls for inoculation quality. Average infection rates for INSV single infection were 96.7% and 93.3% for TSWV single infection (n=30). Therefore, there was no super infection exclusion between TSWV and INSV.

### 3.3. Small RNA (sRNA) profile and distribution reflect INSV antagonism versus TSWV

To look at the relation between the two viruses in mixed infection via profiling of sRNAs, we selected plants inoculated with the ratio that gave us consistent number of successful mixed infection (TSWV: INSV=5:2; w/w). Before sequencing, only plants with O.D. in ELISA tests that were similar for INSV and TSWV in all treatments (single TSWV, single INSV, as well as mixed infections) were used for sRNAs sequencing (2 biological replicates; pooling 3 leaf disks each from 2 infected plants for each biological replicate). ELISA results are based on amount of CP and thus are a good proxy for virions. To measure the ratio of viral RNA in these plants, qPCR was carried out on the RdRp genes from both viruses. Mixed infected plants showed lower Ct values for TSWV, even though they exhibited similar O.D. in ELISA tests for both viruses. The ratio of the viral RdRp were 1:0.56 and 1:0.34; INSV: TSWV for replicate 1 and 2 respectively, indicating an overall lower amount of TSWV than INSV in the samples and the same amount of INSV in both replicates.

Reads that were mapped to either viral or host genome with zero mismatches were used in the analysis. For mock inoculated plants, 60.8% and 62.5% of sRNAs were of host origin (Figure 2). For INSV infected plants, 34.0% and 36.8% sRNAs were of host origin and 34% and 29.4% reads mapped to the INSV genome (Figure 2). Similar trends were observed in TSWV infected samples, but with higher percentage of viral small RNAs (vsRNAs) (47.9% and 43.0% respectively) and lower percentage of host endogenous sRNAs (26.6% and 30.5%, respectively) (Figure 2). VsRNAs from both TSWV and INSV were found in mixed infected samples. The percentages of vsRNAs aligning to TSWV from mixed infections were much less abundant (4.1% and 4.2%) compared to the percentages of TSWV vsRNAs in single infection (47.9% and 43%) and to the percentages of INSV vsRNAs in mixed infection (31.8% and 27.1%) (Figure 2). This reflected the results we found from qPCR of RdRp genes (section 3.1). Host endogenous sRNAs levels of 33.7% and 34.6% in mixed infected plants were in line with the ones found in single infections.

**Figure 2.**
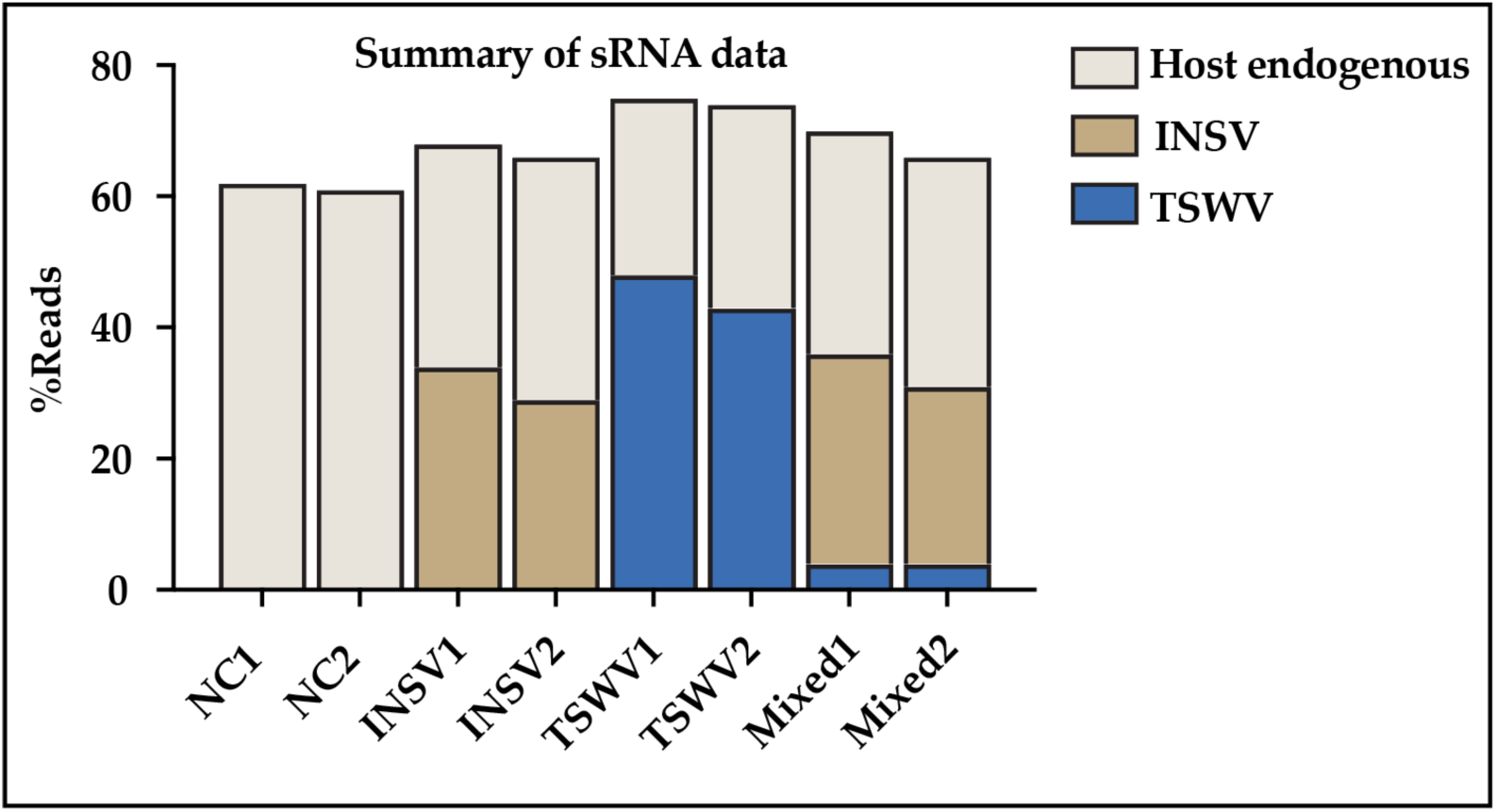
Summary of small RNA reads.

We then mapped and visualized the sRNAs throughout the viral genomes. Our data showed that the pattern of hotspots distribution of INSV vsRNAs on the INSV genome on the positive and negative sense was similar between single and mixed infection (Figure 3a). However, TSWV vsRNA distribution changed dramatically from single to mixed infection, decreasing at the S segment (position 13689-16663) and reaching a high amount at the M segment (position 8915-13679) (Figure 3b). This difference between single and mixed infection only for TSWV, thereby indicates a virus-virus interaction and/or unique and distinctive host-virus interaction. All profiles of sRNAs across all genomes for single and mixed infections and on both polarities showing consistency among biological replicates are in Figure S1 and S2.

**Figure 3.**
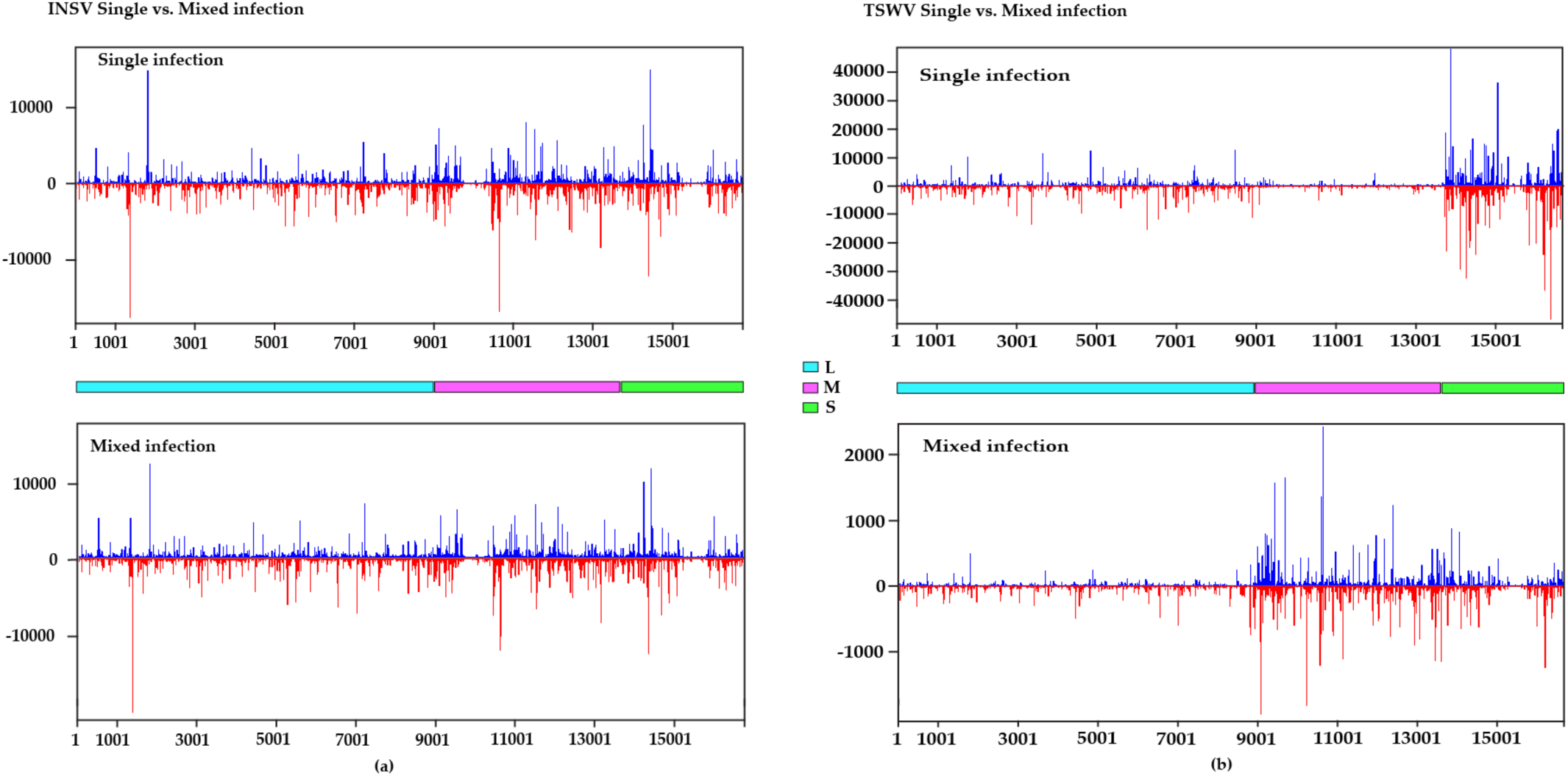
Hotspot distributions on the viral genome. Peaks represent multiple reads aligned to the genome in the same position. Y-axis: number of reads. X-axis: nucleotide position on the viral genome. **(a)** INSV single and mixed infection. **(b)** TSWV single and mixed infection. Blue: viral sense; Red: viral antisense. Figures were generated using MISIS. L segment: 1-8914nt, M segment 8915-13679nt and S segment 13689-16663nt (represented by the consecutive segments with color codes on the solid lines in between top and bottom panel.

Since the molar ratio of each genomic segment during infection is unknown as well as their RNA structure, we chose to report the percentage of vsRNA, as previously done in other studies characterizing the sRNA profiles of TSWV infected plants [38,39]. In those studies, the assumption was that an equal molar concentration of different segments was present during infection, and that the viral RNA lacked secondary structure that would protect it from RNAi. Similarly, to the above-mentioned studies and contrary to the null hypothesis stated above, results from this study showed that vsRNAs alignment was not proportional to the length of the genome segments, moreover, allocation to each of the segments was virus specific (Figure 4a and 4b). The L segments of INSV from single infection accumulated less vsRNAs (on average 12% less than expected, based on the size of the L RNA) (Figure 4a). This may be due to lower accumulation of L RNA in infected plants compared with other segments, as seen by others [46,47]. On the other hand, the M and S segments accumulated more vsRNAs (on average 8.8% and 3.2% more than expected, respectively) (Figure 4a). INSV from mixed infection showed similar vsRNAs accumulation on each genomic segment as INSV from single infection (Figure 4a). However, consistent with the hotspot distribution, TSWV vsRNAs from single infection were very different from the mixed infection (Figure 3b). In single infection, both L and M segments of TSWV accumulated less vsRNAs than expected (with an average of 20% and 22.1% less, respectively), whereas the S segments accumulated much more vsRNAs compared to the expected based on size (on average, 40% more). (Figure 3b). The higher TSWV vsRNAs at NSs region compared to the INSV (an average of 45.5% vs 31.7% were produced in single infections), could suggest that the TSWV isolate might had a weaker silencing suppressor, compared to INSV, while during the mixed infections, the accumulation of TSWV vsRNAs on the same region was still higher than expected by it decreased compared to the single TSWV infection. In a PVX and PVY co-infection, it has been reported that PVX was aided by the silencing suppressor of PVY [48], and it would be interesting to study if tospovirus silencing suppressors could protect each other in mixed infections.

**Figure 4.**
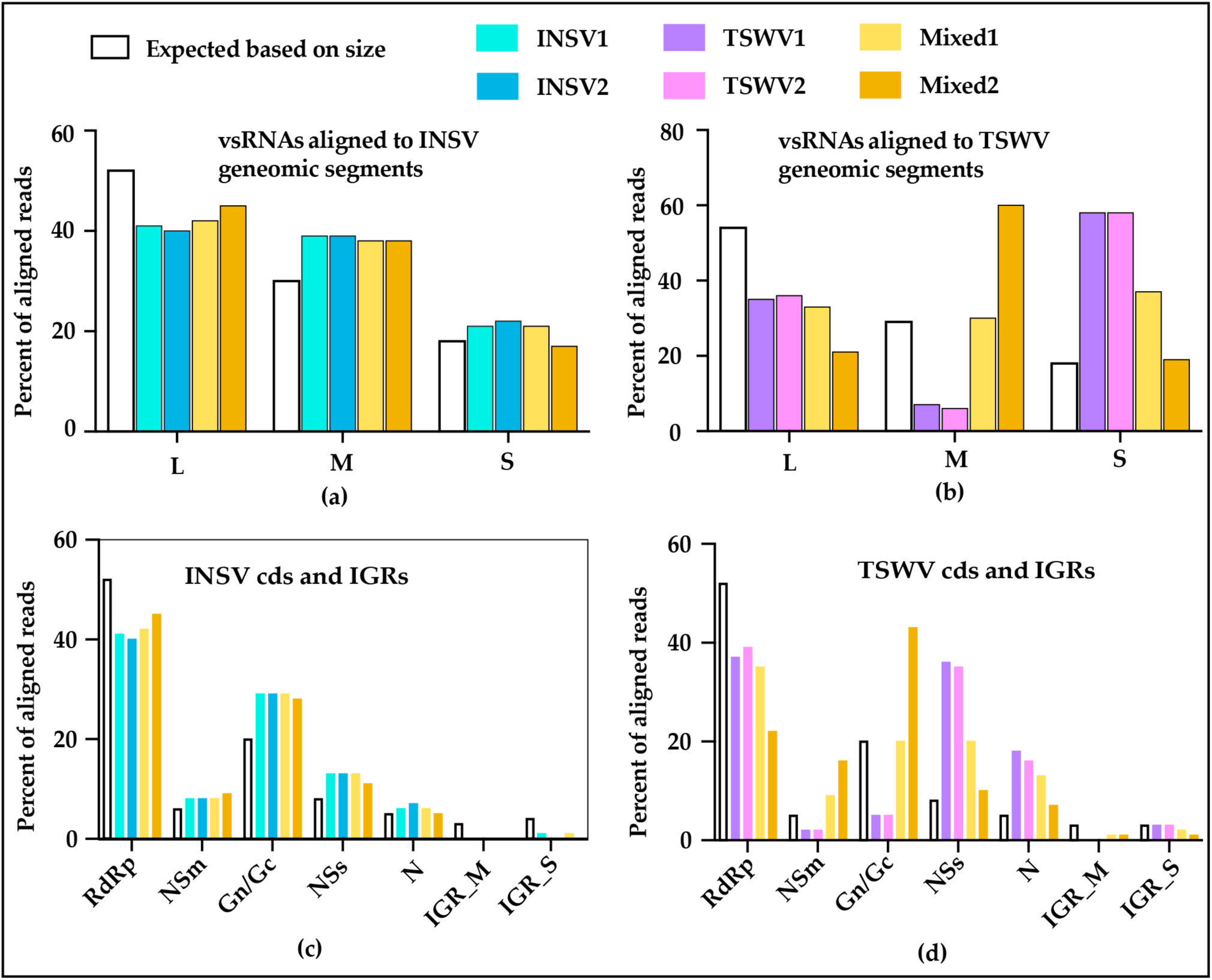
vsRNAs aligned to each segment, cds and IGRs. **(a)** vsRNAs on INSV segments. **(b)** vsRNAs on TSWV segments. **(c)** vsRNAs on INSV cds and IGRs. **(d)** vsRNAs on TSWV cds and IGRs.

The most noticeable difference was noticed in the mixed infection on the TSWV M segment, where accumulation of TSWV vsRNAs got much higher (23-53% higher than single infection) (Figure 3b). The M segment encodes the viral movement protein and the glycoproteins that are involved in genome encapsidation and virus transmission, respectively. Future experiments need be done to study the mechanism behind the differential accumulation of the TSWV M segment during coinfection with INSV, and if these changes correspond to changes in TSWV cell to cell movement, virion formation or transmission. To determine if these differences are constant in the virus coding sequences compared to the intergenic region (IGRs) where regulatory sequences are usually found, we checked the sRNA alignment on the open reading frames (ORF) and IGRs. Percentages of vsRNAs aligned to different ORFs and IGRs of INSV in mixed and single infection were consistent with each other (Figure 4c), as expected based on the data above (Figure 3a, 4a). In case of TSWV, as expected, in single infection, accumulation was higher only at NSs and N (part of S segment), whereas, in mixed infection we found overrepresentation of NSm and Gn/Gc (part of M segment) and NSs and N (much lower than single infection though) (Figure 4d). This pattern was observed in the IGRs of M and S segments as well (Figure 4d).

### 3.4. Different DCLs and AGOs could be recruited during co-infection

sRNAs of 24 nt in length were abundant in mock inoculated plants (Figure 5a), whereas most of the host endogenous sRNA (Figure 5a) and vsRNA in infected samples were 21 and 22nt (Figure 5b, 5c). For INSV infection, 21nt vsRNAs had the highest abundance (Figure 5b), while for TSWV infection, 22 nt vsRNAs had the highest level of accumulation (Figure 5c). VsRNAs of INSV in mixed infected plants showed the same trend as the one seen in INSV single infection (Figure 5b). However, vsRNAs of TSWV in mixed infected plants had a size distribution pattern more like the one seen in INSV infection rather than TSWV infection (Figure 5c). High level of accumulation of 21 and 22 nt vsRNAs has been seen in other virus infected plants and different DCLs were involved during vsRNAs generation [49,50]. In *A. thaliana*, DCL4 is predominantly responsible for the production of 21nt vsRNAs, and DCL2 for production of 22nt vsRNAs [49,50]. Highest abundance of 21nt vsRNA was in fact observed with a TSWV American isolate [38] and an Italian wildtype isolate: p202/3WT [39]. On the contrary, we found 22 nt had the highest level of accumulation in TSWV, which was also reported for the second Italian TSWV isolate considered together with the above mentioned p202/3WT [39], indicating that DCL2 was highly involved in TSWV infection. While the previous studies indicate that the use of DCLs was isolate dependent, our study infers it was species specific. Our results suggest that DCL4 and DCL2 play major role during infections, and that INSV and TSWV elicit different DCLs, when infecting alone, but both DCLs could be involved when plants are infected by both viruses simultaneously.

**Figure 5.**
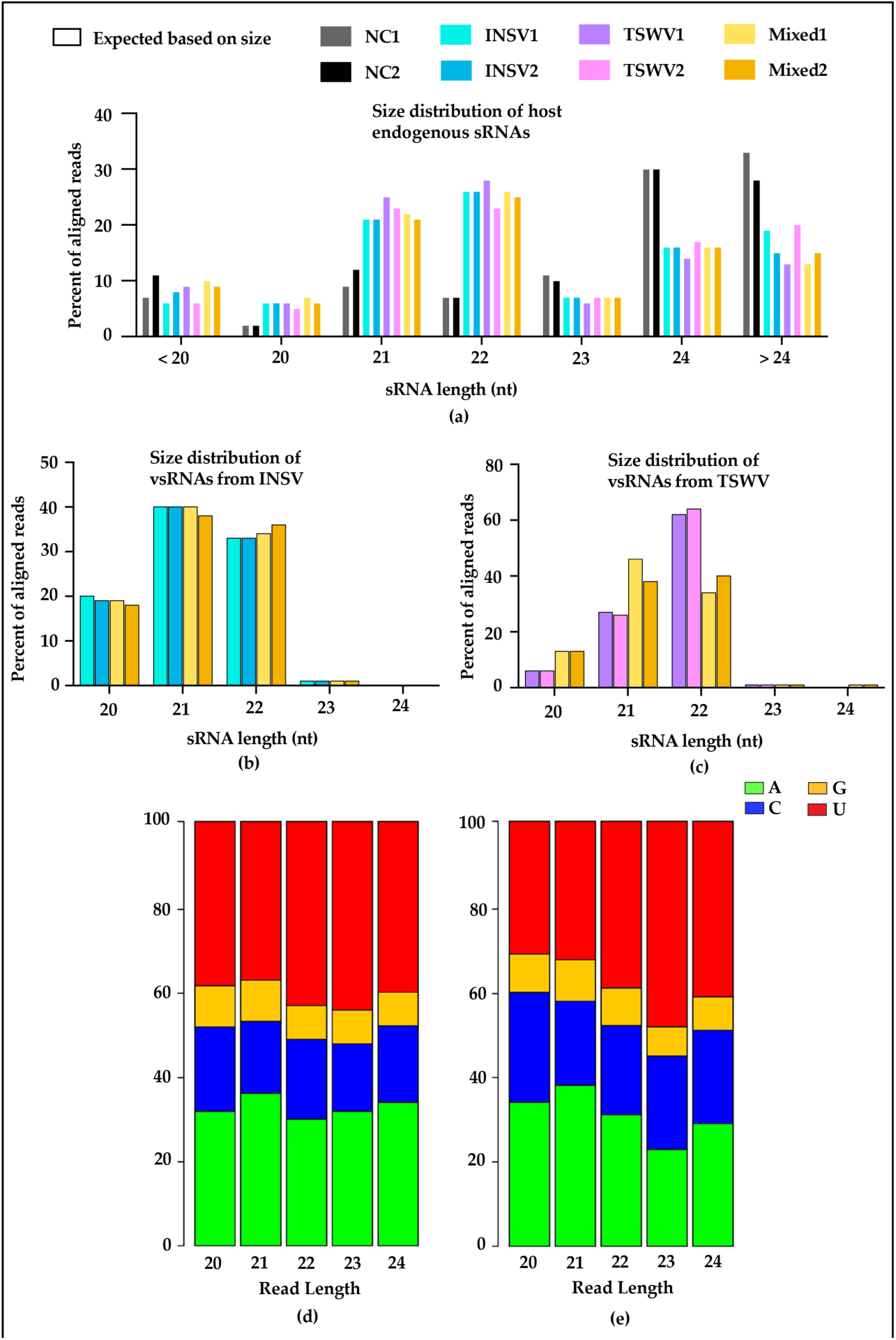
Summary of small RNA reads based on their size. NC: mock-inoculated negative control. Size distribution of host endogenous sRNAs, vsRNAs from INSV, and vsRNAs from TSWV **(a, b, and c)**. Relative frequencies of the 5′ terminal nucleotides from INSV **(d)** and TSWV **(e)** infected plants. Reads were aligned using Bowtie with 0 mismatch to viral genome. Relative frequencies were calculated with aligned reads, and figures were generated by MISIS. Y-axis: percentage of each nucleotide and X-axis: reads length.

Recruitment of vsRNAs proffered by different AGOs can be predicted by the vsRNAs 5’-terminal nucleotide, as AGO2 and AGO4 predominantly recruit sRNAs with a 5’-terminal Adenosine (A), AGO1 favors the harboring of sRNAs, especially the miRNAs that have a 5’-terminal Uridine (U), and AGO5 is biased toward sRNAs with 5’-terminal cytosine (C) [51–54]. This 5’-terminal nucleotide-specific bias is mostly independent of the size of the sRNAs and the biogenesis pathways that synthesize them [51]. In this study, we observed a preference for A and U in the 5’-terminal end (Figure 5d, 5e) in all samples (Figure S3) under both virus treatments, consistent with previous studies on TSWV infected plants [38,39]. This suggests the involvement of AGO1, AGO2, and AGO4 in plant small RNA-mediated defense against TSWV and INSV. Previous studies have also reported that AGO1 and AGO2 are associated with the majority of 21nt sRNAs, while AGO4 is mostly associated with 24nt sRNAs. AGO5, on the other hand, is associated with sRNAs of different lengths (21, 22, or 24nt). As mentioned above, the mock inoculated plants had an abundance of 24 nt sRNAs (Figure 5a), whereas the host endogenous sRNA (Figure 4a) and vsRNA in infected samples were 21nt or 22nt in length (Figure 5b, 5c). This suggests the predominant contribution AGO1 and AGO2 in infected plants and indicates that majority of sRNAs with 5’-terminal A were processed by AGO2 rather than AGO4. The increase of sRNA with 22nt in length, especially in the plants with single TSWV infection also indicates the possible involvement of AGO5 in TSWV infection.

### 3.5. Virus-activated small interfering RNAs (vasiRNAs) in virus infected plants

During a virus infection, virus-activated small interfering RNAs (vasiRNAs) are synthesized from host-gene transcripts and differ from the other endogenous sRNAs of host origin. VasiRNAs, mostly 21 and 22nt in length incorporated into AGO1 and AGO2/RISC, cause silencing of target mRNAs of host genes [55]. vasiRNAs were first described in 2014 [55] where *A. thaliana* infected with silencing suppressor-deficient viruses accumulated endogenous sRNAs predominantly 21nt in length. In our experiment, both mock inoculated controls showed the highest abundance of 24nt sRNAs, which is the typical size of host endogenous sRNAs, and a lower abundance of 21, 22 and 23nt sRNAs (Figure 5A). In contrast, most host endogenous sRNA in infected samples were 21nt or 22nt in length (Figure 5A), which indicates the possibility of vasiRNAs production in the infected plants. In *A. thaliana* DL4/RDR1, AVI2H and AGO2 are required for vasiRNA biogenesis [55], however, nothing is known about this same process in *N. benthamiana* and mutants for these enzymes should be used to characterize this process in this host.

### 3.6. MiRNAs vary in INSV and TSWV infection and are more similar in INSV and mixed infections

The involvement of AGO1 in the biogenesis of miRNAs prompted us to investigate miRNA production and their differential regulation in infected plants. A principal component analysis (PCA) plot based on the *miRNA* loci demonstrated a strong clustering of biological replicates from mock-inoculated (NC) and mixed infected treatments compared to those from INSV and TSWV treatments (Figure 6a). Our data exhibited, out of 174 *miRNA* loci identified across all treatments, 80 (20 belonging to known miRNA families and 60 unknown) were significantly differentially regulated (based on FDR ≤ 0.1, true difference > 2-fold after Benjamini–Hochberg correction for multiple testing) in at least one comparison with most of them being up-regulated, and a few down-regulated during infection (Figure 6, Table S1). The number and type of miRNA differentially regulated in each comparison varied, except for the mixed infection vs. INSV where the regulated miRNAs remain the same (Figure 6e), suggesting that INSV is the main driver of miRNA regulation in mixed infection. In addition, almost all the 51 *miRNA* loci differentially regulated in mixed infections vs. mock infected controls were also differentially regulated in TSWV and INSV single infections in the same trend (i.e., up, or downregulated), suggesting that miRNAs regulated during orthotospovirus mixed infections are not novel, compared to the combined single infections.

**Figure 6.**
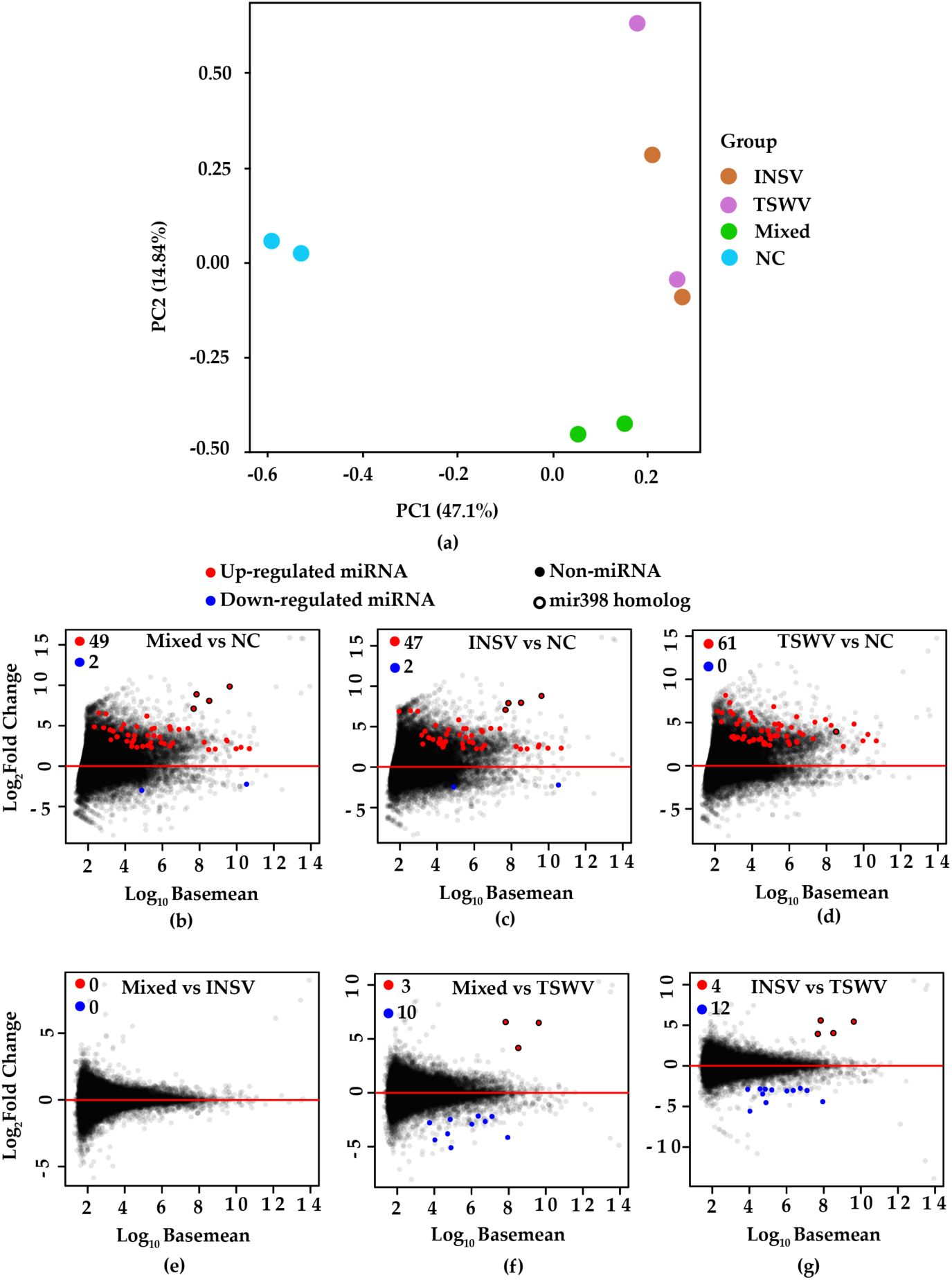
Host miRNAs and their differential regulation in different treatments (mock inoculated control: NC, INSV, TSWV and both infected: INSV, TSWV, and Mixed). **(a)** PCA plot based on de-novo identified host miRNA clusters. **(b-g)** Mean abundance plot of *N. benthamiana miRNA* loci comparing two different treatments. Significantly up-regulated and down-regulated miRNA loci are highlighted (alternative hypothesis: FDR ≤ 0.1, true difference > 2-fold after Benjamini–Hochberg correction for multiple testing).

Compared to mock-inoculated controls, INSV infection (Figure 6c) differentially regulated fewer *miRNAs* than TSWV infection (Figure 6d) (49 vs. 61 loci). Pairwise comparisons between the infected treatments exhibited even fewer differences. For instance, mixed infection vs. INSV and mixed vs. TSWV showed differential regulation of 0 and 13 *miRNA* loci (3 up, 10 down) respectively (Figure 6e, 6f), and INSV vs. TSWV showed 16 loci (4 up, 12 down) (Figure 6g). These results are consistent with the notion that larger differential regulation of miRNAs occurs in plants upon infection, and further changes in infection will result in a lower level of differential regulation.

Among the known miRNAs, the miR398 family was significantly upregulated in all comparisons (Figure 6b, 6c, 6e, 6f, 6g) except for the mixed infection vs. INSV (Figure 6e). miR398 is involved in oxidative stress responses in plants [56] and has been shown to be upregulated in tomato plants during southern tomato virus, a persistent virus [57], and rice stripe virus (an acute virus) infection [58]. In addition, *miR170* and *miR395* were highly upregulated in TSWV infected plants (Table S1), compared to mock, INSV and mixed infected plants. *miR170* is known to be downregulated during tomato leaf curl new delhi virus [59] and turnip crinkle virus infection [60]. The miR395 family is predicted to target the ATP sulfurylase encoding mRNA that is involved in sulfate assimilation, and its expression increases during sulfate starvation [61]. The abundance of this miRNA was also seen to increase by turnip crinkle virus infection [60].

In one of the few studies where miRNAs were analyzed during mixed infections, miRNA accumulation was changed by mixed infection of the potyviruses PVY or plum pox virus (PPV) with the potexvirus PVX. More severe symptoms were produced during the PVY/ or PPV/PVX mixed infection with altered accumulation of miRNA156, miRNA168, miRNA171 and miRNA398. Consistently with the study above, miRNA398 was differentially regulated in INSV and TSWV mixed infection. The similarity in the regulation of these miRNAs during single and mixed plant infection suggests that some plant responses are conserved during orthotospovirus infection (Figure 6). Furthermore, our study shows that INSV seems to play a prominent role during plant regulation in single as well as mixed infections.

## Conclusions

This study demonstrates that INSV and TSWV have an antagonistic relationship, and INSV possesses better fitness in *N. benthamiana* during mixed infection than TSWV, resulting in a higher incidence of infection and a higher titer than TSWV and higher amount of INSV svRNAs.

These findings support results seen in Zhao and Rosa, 2020, where thrips were shown to preferentially transmit INSV from TSWV and INSV mixed infected plants [26]. While the exact mechanism explaining this antagonistic relationship is unclear, our study provides insights about varied host responses involving RNAi, to single and mixed orthotospovirus infection. Interestingly, while INSV vsRNA and miRNA accumulation didn’t change much between single and mixed infections, TSWV showed marked differences during mixed infections.

Our study indicates that while plants use conserved mechanisms to act against viruses, differences must exist for where or how orthotospoviruses interact with the plant RNAi machinery. These differences and similarities could be exploited to generate novel orthotospovirus species or genus specific management strategies.

## Supplementary Materials

The following supporting information can be downloaded at: www.mdpi.com/xxx/s1, Figure S1: Hotspot distributions on the viral genome. a) INSV vsRNAs from INSV1, b) INSV vsRNAs from INSV2, c) INSV vsRNAs from Mixed1, d) INSV vsRNAs from Mixed2. Blue: viral sense; Red: viral antisense Figures were generated using MISIS. L segment: 1-8774nt, M segment 8775-13751 nt and S segment 13752-16761nt.; **Figure S2:** Hotspot distributions on the viral genome. a) TSWV vsRNAs from TSWV1, b) TSWV vsRNAs from TSWV2, c) TSWV vsRNAs from Mixed1, d) TSWV vsRNAs from Mixed2; Blue: viral sense; Red: viral antisense. Figures were generated using MISIS. L segment: 1-8914nt, M segment 8915-13679 nt and S segment 13689-16663nt.; **Figure S3:** Relative frequencies of the 5’ terminal nucleotides a) INSV vsRNAs from INSV1, b) INSV vsRNAs from INSV2, c) INSV vsRNAs from Mixed1, d) INSV vsRNAs from Mixed2, e) TSWV vsRNAs from TSWV1, f) TSWV vsRNAs from TSWV 2, g) TSWV vsRNAs from Mixed1, h) TSWV vsRNAs from Mixed2. Green: A, blue: C, yellow: G and red: U. Reads were aligned using Bowtie with 0 mismatch to viral genome. Relative frequencies were calculated with aligned reads, and figures were generated by MISIS. **Table S1:** Differential expression of miRNAs across treatments.

## Author Contributions

Conceptualization, K.Z. and C.R.; investigation, K.Z. and N.R.J; writing— original draft, K.Z., M.T.I. and C.R.; writing—review and editing, K.Z., M.T.I., N.R.J., M.J.A. and C.R; visualization, K.Z., M.T.I, and N.R.J.; formal analysis, K.Z., M.T.I, and N.R.J.; funding acquisition, C.R. All authors read and agreed to the published version of the manuscript.

## Data Availability Statement

The datasets generated and analyzed for this study can be find in the NCBI with Accession Number: PRJNA833054.

## Acknowledgments

The authors would like to thank Dr. Rich Marini for his assistance on experimental design for virus-virus interaction, Dr. Feng Qu for his advice on super infection exclusion study and Dr. Craig Praul for his assistance with small RNA sequencing, Chauncy Hinshaw for his help with sequencing data deposition.

